# The neural basis of swap errors in working memory

**DOI:** 10.1101/2023.10.09.561584

**Authors:** Matteo Alleman, Matthew Panichello, Timothy J. Buschman, W. Jeffrey Johnston

## Abstract

When making decisions in a cluttered world, humans and other animals often have to hold multiple items in memory at once – such as the different items on a shopping list. Psychophysical experiments in humans and other animals have shown remembered stimuli can sometimes become confused, with participants reporting chimeric stimuli composed of features from different stimuli. In particular, subjects will often make “swap errors” where they misattribute a feature from one object as belonging to another object. While swap errors have been described behaviorally, their neural mechanisms are unknown. Here, we elucidate these neural mechanisms through trial-by-trial analysis of neural population recordings from posterior and frontal brain regions while monkeys perform two multi-stimulus working memory tasks. In these tasks, monkeys were cued to report the color of an item that either was previously shown at a corresponding location (requiring selection from working memory) or will be shown at the corresponding location (requiring attention to a position). Animals made swap errors in both tasks. In the neural data, we find evidence that the neural correlates of swap errors emerged when correctly remembered information is selected incorrectly from working memory. This led to a representation of the distractor color as if it were the target color, underlying the eventual swap error. We did not find consistent evidence that swap errors arose from misinterpretation of the cue or errors during encoding or storage in working memory. These results suggest an alternative to established views on the neural origins of swap errors, and highlight selection from and manipulation in working memory as crucial – yet surprisingly brittle – neural processes.

## 1 Main

Whether recalling items on a shopping list, or matching new names to faces, reckoning with the limits of one’s working memory is an inescapable part of human experience. Behavioral studies have characterised limits in the storage and selection of multiple items from working memory [1–5]. For instance, when subjects are cued to report one of the features (e.g., the color) of a target item from a set of remembered items, the likelihood of an error is increased by either increasing the number of items that must be remembered[3, 4, 6] or decreasing the time for which the items are presented[7, 8].

Previous work has provided insight into the neural mechanisms that lead to the drift of memories over time and the ‘forgetting’ of items[9, 10], but a third type of error – the ‘swap error’ – is not well understood. Swap errors occur when a subject reports a feature value that was present among the remembered items but that does not correspond to the target item[4, 11]. For example, if a red square and blue circle are shown to a subject, when asked what color the square had been the subject might make a swap error and report ‘blue’.

Swap errors have been of particular interest not only because they provide insight into fundamental limitations of working memory[12, 13], but also because they may arise from failures to correctly bind the distinct features of a particular stimulus together[14–18]. In the previous example, the subject might say the square was blue because they actually remember seeing a blue square and red circle. However, the relationship of swap errors to binding has been controversial, as different explanations – such as an error when interpreting or representing the cue[19] or incomplete encoding of the stimulus array[20] – can give rise to the same behavioral pattern. For instance, the subject might mishear the experimenter and say blue because they thought they were supposed to report the color of the circle. While recent work has shown that swap errors are reflected in neural activity after the target is known but before the response period[21] – and thus do not reflect a last minute guess – existing work leaves many possible explanations for swap errors open.

To address the controversy around the nature of swap errors, we analyzed neural population recordings from macaque monkeys performing a color working memory task. Consistent with previous work in humans, on a subset of trials monkeys made swap errors by incorrectly reporting the color of the distractor (rather than the target). Surprisingly, we find evidence for a novel explanation of swap errors: they occurred during the selection of correctly remembered information from working memory. We replicated this pattern of results in a second attention task, which also elicited swap errors and had a similar neural mechanism. Overall, this work provides new insight into the neural mechanisms underlying widely observed limitations in working memory – and highlights the fragility of manipulating information in working memory.

Monkeys performed two versions of a continuous working memory task (fig. 1a-b)[22]. In the ‘retrospective’ task, the animal remembered two colors, one presented at an upper spatial position (the upper color) and one presented at a lower spatial position (the lower color; fig. 1a). After a memory delay period, an abstract cue reliably indicated whether the color at the upper or lower position was the ‘target’. After a second memory delay, the animal reported the color of the target by making a saccade to the matching color on a color wheel. Note, the color wheel was rotated on every trial to prevent motor planning. Therefore, the retrospective task required the animal to hold two color-position pairs in working memory before selecting one, and reporting the corresponding color. Monkeys also performed a ‘prospective’ task which was identical with the exception that the cue was shown before the colors (fig. 1b). In this condition, the monkey could use visual attention to select the relevant target stimulus immediately and ignore the other ‘distractor’ stimulus[23, 24]. Previous work with this task has shown that the neural representations of the items in working memory are distributed across prefrontal, parietal, and visual cortex in both tasks[22]. It also showed that the retrospective selection and prospective attention conditions were surprisingly similar: In both cases, the representation of the target stimulus was dynamically transformed – in the same way – to facilitate the animal’s behavioral response[22].

**Figure 1:**
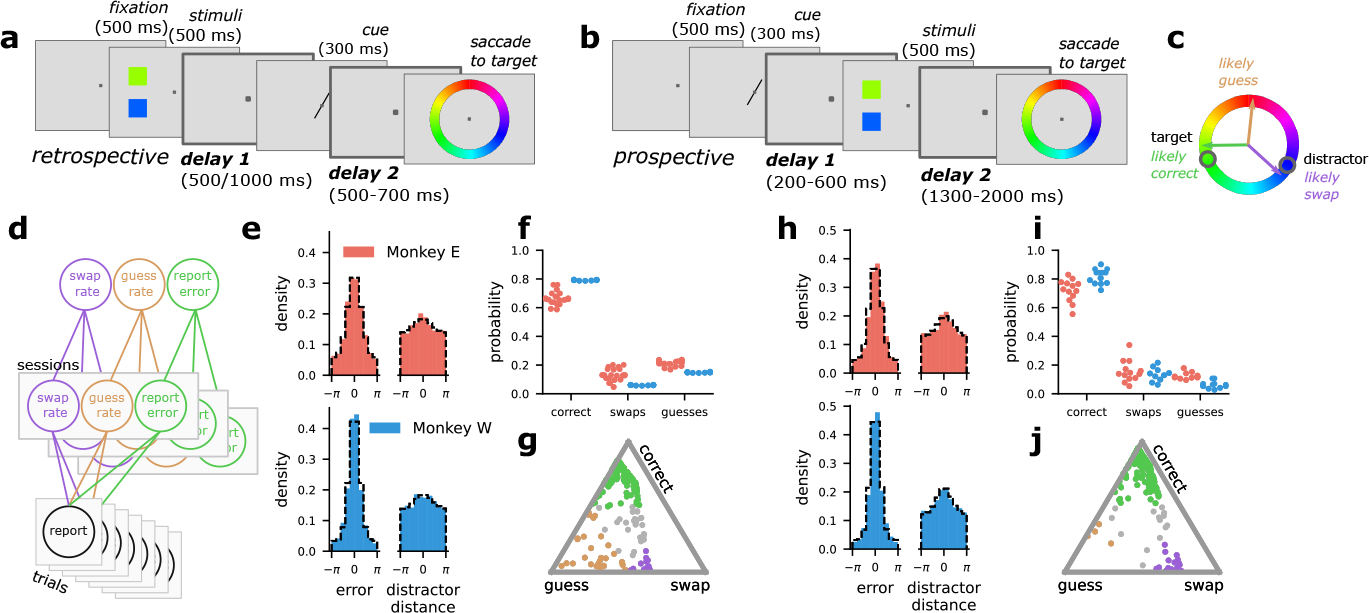
The retrospectively and prospectively cued working memory tasks, as well as the behavioral modeling approach. **a** Schematic of the retrospective task. **b** Schematic of the prospective task, note that the order of the cue and stimuli is switched with respect to the respective task shown in **a. c** Schematic of the possible trial outcomes: correct response (close to the target), a guess (uniform random), and a swap (close to the distractor). **d** Schematic of the behavioral modeling approach. Each trial response is explained in terms of three parameters: the rate of swap errors, the rate of guesses, and the dispersion around the target response. A single model is fit hierarchically across all the sessions, where the session level parameters (middle) are modeled as being sampled from a distribution of animal-level parameters (top). The model is fit separately for each animal. **e** Histograms of the errors from the target (left) and the distractor stimulus (right) for Monkey E (top) and Monkey W (bottom) on retrospective trials. The dashed black line is the posterior predictive distribution from the behavioral model. The concentration around zero for the plots on the right indicate that both monkeys make swap errors. **f** The inferred rate of each response type on retrospective trials for each session (points) and both monkeys (colors). **g** An example session from Monkey E on the retrospective task. Each point is the response on a particular trial. The triangle is the three-dimensional simplex for the probabilities of each response type. Each of the three corners represent probability 1 of a particular response type. Colored points have probability *>* .5 of belonging to their respective response type. **h - j** The same as **e - g** but for prospective trials, and the example session in **j** is from Monkey W.

In the tasks, the animals made three qualitatively different kinds of responses (fig. 1c): correct responses; guess responses, where the animal chooses a random color; and swap responses, where, as introduced above, the animal reports the color of the wrong (distractor) memory. Following the human working memory literature[4] (but see [25]), we model the distribution of the behavioral responses within each task (fig. 1e, h) as arising from a mixture model with probabilities for each response type (fig. 1d, and see *The behavioral model* in *Methods*). The generative model produced by the fit closely matched the empirical distribution of animal responses (fig. 1e, h, black dashed-line compared to colored histogram).

This model shows significant evidence for all three types of response across both the retrospective and prospective tasks. In the retrospective task, both monkeys performed the majority of trials correctly (Monkey E: correct probability = 0.60 to 0.72, Monkey W: correct probability = 0.76 to 0.82, unless otherwise noted all reported ranges are 95 % confidence intervals for the relevant quantity; fig. 1e, f) but also made guess (Monkey E: guess probability = 0.16 to 0.27, Monkey W: guess probability = 0.12 to 0.18) and swap errors (Monkey E: swap probability = 0.08 to 0.19; Monkey W: swap probability = 0.04 to 0.08). There was a similar pattern in the prospective task (fig. 1h, i, Monkey E: correct probability = 0.65 to 0.78, guess probability = 0.09 to 0.18, swap probability = 0.10 to 0.22,; Monkey W: correct probability = 0.73 to 0.85, guess probability = 0.03 to 0.13, swap probability = 0.09 to 0.19,). Using the fitted model, we computed the posterior probability that a particular trial arose from each response type (fig. 1g, j, bottom), which we incorporated into our probabilistic model of the neural population data below.

We recorded single and multi-unit neural activity simultaneously from posterior parietal cortex, frontal cortex, motor cortex, posterior inferotemporal cortex, and visual area 4 (posterior parietal cortex: 10 to 39 units; motor cortex: 18 to 53 units; prefrontal cortex: 22 to 57 units; V4 and posterior IT: 15 to 33 units; frontal eye fields: 2 to 23 units, where the ranges give the minimum and maximum number of simultaneously recorded neurons across sessions). Since swap errors are rare and each specific trial configuration was also rare, we were unable to combine the neural populations across sessions. Thus, to increase our statistical power, we took advantage of the distributed nature of working memory representations[22, 26] and analyzed all of the units recorded in a single session as a single population, combining across the different brain regions (combined: 80 to 181 units). We performed a region exclusion analysis to determine if the observed effects depended only on neural activity from specific regions; this analysis found no consistent effects across monkeys (see *Region-dropping analysis* in *Supplement* and fig. S1).

We then compared the neural activity on trials with correct or swap responses to the neural representation that would be expected if the color of each item were faithfully represented (“nominal rep.”, fig. 2a, gray star) and several alternate neural representation (e.g., the right colors represented in the wrong positions; fig. 2a, purple star). We begin by analyzing the first delay period of the retrospective task (fig. 2a). In this period, the monkey must remember both the two colors and their locations; so, we expect that both colors will be represented in the neural population activity. In particular, we fit the neural activity with the linear model,

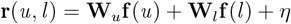

where **W**_*u*_ and **W**_*l*_ are fitted matrices, *u* and *l* are the colors presented in the upper and lower positions, respectively, **f** (*·*) is a function that transforms the color (given in radians) to a representation in spline basis functions (see *The neural mixture model* in *Methods*), and *η* is noise. Given a green upper color and a blue lower color, the nominal neural representation is r_nominal_ = **r**(green, blue). One alternate neural representation that we consider here is a misbound representation, r_misbinding_ = **r**(blue, green), where color and position are mis-associated. That is, if behavioral swap errors arise from a misbinding between color and spatial position, then we expect the neural representation on trials with behavioral swap errors to resemble the misbound representation r_misbinding_ (fig. 2a, “misbound rep”) rather than the nominal representation r_nominal_. On trials with correct behavioral responses, we expect that the neural representation will resemble the nominal representation (fig. 2a, “nominal rep.”).

**Figure 2:**
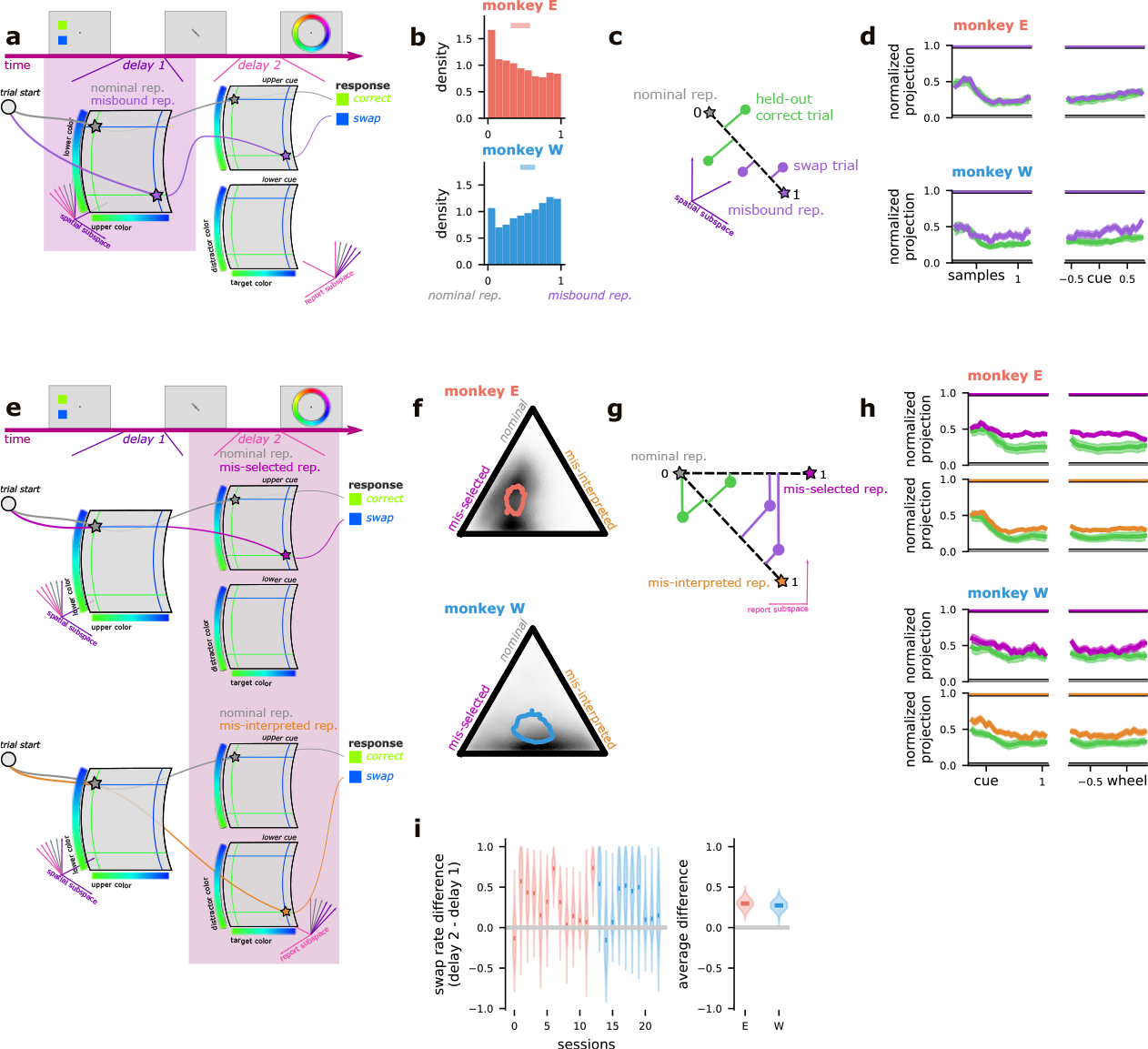
The neural correlates of swap errors in the retrospectively cued working memory task. **a** Schematic of an example trial during the first delay period of the retrospective task. The nominal (gray star) and misbound (purple star) representations are shown for this example trial. The misbound representation manifests as a swapped representation of the upper and lower colors (green-upper, blue-lower to blue-upper, green-lower). **b** The average (top bar) and aggregate posterior (distribution) across sessions for the p_misbind_ parameter in the neural mixture model for the first delay period. Values close to one indicate an association between swap errors and misbound representations, values close to zero indicate that trials with swap errors are still associated with the nominal representation. **c** Schematic of the cross-validated version of the analysis. Nominal and misbound representations (stars) are constructed for each trial using the same kind of linear model as used in the neural mixture model. The model is fit only on likely correct trials. Then, held-out correct trials and swap trials (circles) are projected onto the dimension connecting the nominal and misbound representations (dashed line). The distance is normalized to be between zero and one. **d** Time course of evidence for an association between swap errors and the misbound representation under the analysis in **c**, for each monkey (top and bottom) as well as centered on the time of sample (left) and cue (right) presentation. **e** Schematic of the same example trial from above, showing the contrast between the nominal representation (gray stars) and the mis-selected colors representation (top, pink star) as well as the mis-interpreted cue representation (bottom, orange star). **f** The average (contour outline) and aggregate posterior (heatmap) evidence for the nominal, mis-selected colors, and mis-interpreted cue representations under the neural mixture model in delay 2. **g** Schematic of the cross-validated version of the analysis. Correct, color selection, and cue interpretation prototypes are constructed using a linear model fit on likely correct trials. Then, held-out correct trials and likely swap trials are projected along the axes connecting the correct prototype with each of the error prototypes and normalized as before. **h** Time course of evidence for the mis-selected colors and mis-interpreted cue representations for both monkeys (top and bottom) as well as centered on the cue presentation (left) and appearance of the response wheel (right). **i** The difference between the evidence for an association between swap errors and the alternative representations in delay 1 and delay 2 under the neural mixture model for each session (left) and averaged across all sessions (right; positive values indicate more evidence for correlates in delay 2).

We express our hypotheses about neural representations in a mixture model, with a parameter controlling the probability that the neural activity is sampled from a distribution around the misbound representation instead of the nominal representation. This approach allows us to avoid a hard categorization of trials into correct and swap responses. Instead, we use the posterior probability that a trial had a correct or swap response under the full behavioral model. For an observed response r_observed*i*_ from a particular trial *i*, the mixture model has the following form,

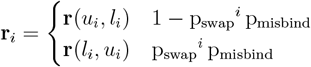

where p_swap_ is the posterior probability that the given trial has a behavioral swap response (as inferred by the behavioral model, fig. 1g) and p_misbind_ is a fit parameter (see *The neural mixture model* in *Methods* for more details). Note that the observed, nominal, and misbound responses as well as the probability of a behavioral swap response are all indexed by trial while the probability of a misbinding error p_misbind_ is not. Intuitively, if p_misbind_ = 1, then this means that all of the trials with behavioral swap responses also have neural representations that resemble the misbound representation. If p_misbind_ = 0, then this means that all of the trials with behavioral swap responses have neural representations that resemble the nominal representation. We also include a hypothesized explanation for guess trials in the full neural mixture model (see *Analysis of guess responses* in *Supplement* and fig. S2).

We do not find strong evidence that behavioral swap errors are associated with misbound neural representations in either monkey (fig. 2b); the aggregate posterior distribution of p_misbind_ is skewed toward 0 in one monkey (Monkey E: 0.30 to 0.58 probability of alternate rep. on swap trials; fig. 2c, top) and toward 1 in the other (Monkey W: 0.44 to 0.64; fig. 2c, bottom). In both monkeys, however, only a handful of individual sessions have posterior distributions that strongly favor an association between swap errors and misbound representations (Monkey E: significantly greater than 0.25 in 3/13 sessions; Monkey W: 2/10 sessions).

We perform a second, fully cross-validated, sliding window, version of this analysis to confirm these findings across time (fig. 2c and see *Cross-validated neural model* in *Methods* for more details). Here, we used correct trials to fit the linear model described above. Then, for each held-out correct and swap trial, we used the linear model to predict corresponding nominal and misbound representations (fig. 2c, green and purple stars, respectively). These pairs of predicted representations defined a single dimension in population space for each trial, onto which we projected the activity from that trial (fig. 2c, green and purple circles for correct and swap trials, respectively). If swap errors are associated with misbound representations, then we expect swap trials to be closer – on average – to the misbound representation than correct trials are.

The cross-validated analysis finds no difference between correct and swap trials in the second monkey just after the stimuli appear (0 ms to 500 ms after the stimuli appear, Monkey W: activity on swap trials is − 0.07 to 0.07 closer to the misbound rep. than on correct trials; fig. 2d, bottom left), but does find a significant difference prior to the appearance of the cue (− 500 ms to 0 ms before the cue appears, Monkey W: 0.01 to 0.17; fig. 2d, bottom right). These results suggest that some swap errors may emerge while multiple stimuli are remembered across a delay period, consistent with some previous theories[15, 16]. However, there is no significant difference between swap and correct trials in the first monkey in either time period (0 ms to 500 ms after the stimuli appear, Monkey E: − 0.09 to 0.01; − 500 ms to 0 ms before the cue appears, Monkey E: − 0.03 to 0.08; fig. 2d, top row). Overall, our results suggest that some behavioral swap errors in this task may be due to neural misbinding, but the effect is subject dependent.

Next, we perform a similar analysis during the second delay period of the retrospective task. At this point of the task, the monkeys have seen a cue that indicated which of the two items to select and, eventually, report. Therefore, we incorporate a binary “cue” variable, *c*, into our model:

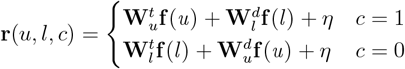

where we now have four parameter matrices, one each for the upper and lower color representations when they are the target or distractor. For example, 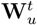 is the representation of the upper color when it is the target, while 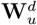 is its representation when it is the distractor.

In the second delay period we hypothesize that behavioral swap responses will be associated with either 1) a mis-selection of the representation of the target or 2) a mis-interpretation of the cue. The mis-selected color representation has a correct representation of the cue, but the actual target color is represented as the distractor and the actual distractor is represented as the target (fig. 2e, top, “mis-selected”). Expressed with our linear model, this would be **r**(*l, u, c*). This representation is consistent with an unreliable transformation from the upper-lower color representations in the first delay period to the target-distractor color representations of the second delay period. These upper-lower and target-distractor subspaces have been shown to be orthogonal to each other[22]. The mis-interpreted cue representation has an incorrect representation of the cue and a representation of the colors that correctly follows the mistaken cue (fig. 2e, bottom, “mis-interpreted”). In our model, this is **r**(*u, l*, 1 − *c*). It is consistent with a mis-interpretation of the meaning of the cue that leads to a reliable, but wrong, transformation between the upper-lower and target-distractor color representations. Finally, correct responses were expected to match the nominal representations that faithfully reflect the target and distractor colors, as well as the identity of the cue (fig. 2e, “nominal rep.”).

As before, we incorporated the nominal and two possible swap representations into a mixture model. In both monkeys the mixture model found strong evidence for neural correlates of swap errors (i.e., trials with a swap error have representations that resemble one of the two alternate representations rather than the nominal representation, fig. 2f; Monkey E: 0.67 to 0.79 probability of alternate rep. on swap trials; Monkey W: 0.74 to 0.88). In addition, every individual session also shows strong correlates of swap errors (Monkey E: significantly greater than 0.25 in 12/13 sessions; Monkey W: 9/10 sessions). In one monkey, the mis-selected color representation is more associated with swap errors than the mis-interpreted cue representation (Monkey E: 0.17 to 0.30 greater probability of mis-selected colors than mis-interpreted cue representation; fig. 2e, top). In the second monkey, there is an even split between the two alternate representations (Monkey W: − 0.20 to 0.16 greater probability of mis-selected colors than mis-interpreted cue representation; fig. 2e), which may be due to a weaker overall encoding of the cue during the second delay period (0.06 to 0.13 higher cue decoding performance in Monkey E than W). Further, using the fully cross-validated timecourse analysis above, we show that the evidence for a mis-selected colors representation on trials with swap errors emerges as the cue is presented in both monkeys (0 ms to 500 ms after the cue comes on, Monkey E: activity on swap trials is 0.08 to 0.20 closer to the mis-selected rep. than on correct trials; Monkey W: 0.05 to 0.23; fig. 2h, left).

Next, we compare the strength of the evidence for errors in delay 1 relative to delay 2. In both monkeys, we find stronger evidence for the neural correlates of swap errors in the second delay period (Monkey E: 0.17 to 0.43 greater in delay 2 than delay 1, Monkey W: 0.14 to 0.40, fig. 2i). Overall, our evidence suggests that swap errors primarily occur due to an erroneous transformation between the location-specific (upper or lower) representation and the target-distractor representation. We refer to this as a selection error. We find only inconsistent evidence that swap errors emerge due to an initial misbinding of color to space or a misinterpretation of the meaning of the cue.

Finally, we analyzed the neural population activity during the two delay periods of the prospective task. First, we investigated whether behavioral swap errors can be explained as a mis-interpretation of the cue during the first delay period (fig. 3a, “mis-interpreted rep.”). To do this, we trained a decoder to distinguish between trials with either an upper-or lower-cue, using only trials with likely correct responses. Then, we tested whether or not this decoder successfully generalizes to trials with behavioral swap responses (fig. 3b). If the decoder fails to generalize, then this would indicate that the cue is mis-remembered or mis-interpreted on swap trials (similar to the hypothesis discussed above). In one monkey, we find that the decoder performs just as well on swap trials as on held-out correct trials (Monkey E: − 0.08 to 0.01 difference in decoding performance, swap - correct, fig. 3c). In the second monkey, there is a small, but significant decrease in decoder performance between held-out correct trials and swap trials (Monkey W: − 0.13 to − 0.03, fig. 3c). Overall, we find inconsistent evidence for cue interpretation errors in the first delay period (and see fig. S4a for this analysis over time).

**Figure 3:**
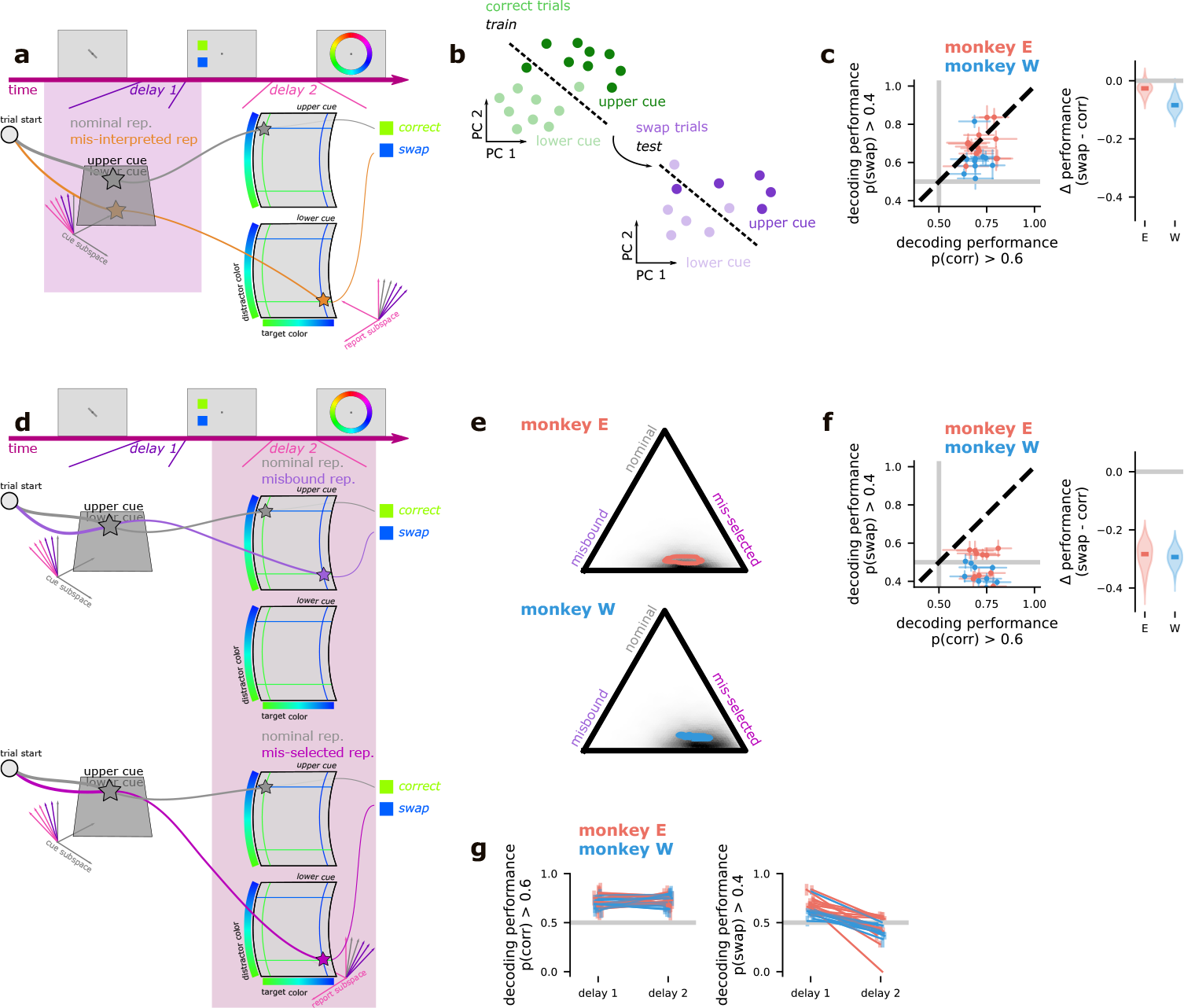
The neural correlates of swap errors in the prospectively cued working memory task. **a** Schematic of an example trial in the prospective task, showing the nominal (gray star) and mis-interpreted cue (orange star) representations. **b** Schematic of the decoder generalization analysis used in **c** and **f**. A linear decoder is trained to discriminate between trials with an upper or lower cue using only likely correct trials (left). Then, that same decoder is tested on likely swap trials (right). **c** The performance of a decoder trained and tested during delay 1 on likely correct trials (left x-axis) and trained on likely correct but tested on likely swap trials (left y-axis), for each session (points) and the difference between the two averaged across sessions (right). **d** Schematic of an example trial in the prospective task, showing the nominal as well as the mis-bound color (top, purple star) and mis-selected cue (bottom, pink star) representations. **e** The average (contour outline) and aggregate posterior evidence for the association of the nominal, misbound, and mis-selected representations with swap errors under the neural mixture model. **f** The analysis schematized in **b** applied to the second delay period. **g** The same decoding approach schematized in **b** and applied to delay 1 and 2 in **c** and **f**, respectively. Here, we compare across delay periods for likely correct trials (left) and likely swap trials (right).

In the second delay period, we can use the same neural mixture model to quantify the likelihood of different alternative representations. This analysis shows strong evidence for neural correlates of swap errors (Monkey E: 0.91 to 0.93 probability of alternate rep. on swap trials; Monkey W: 0.89 to 0.91; fig. 3e). In addition, almost every individual session also shows strong correlates of swap errors (Monkey E: significantly greater than 0.25 in 13/13 sessions; Monkey W: 10/10 sessions). Further, it shows that swap errors in the prospective task are associated with mis-selected cue representations rather than misbound color representations (Monkey E: 0.12 to 0.36 greater probability of mis-selected cue than misbound colors representation, Monkey W: 0.27 to 0.50; fig. 3e). Again, we apply the cross-validated version of the analysis across time and show that the mis-selected cue representation becomes associated with swap errors as the color samples are presented (0 ms to 500 ms after the samples come on, Monkey E: activity on swap trials is 0.16 to 0.24 closer to the mis-selected rep. than on correct trials; Monkey W: 0.23 to 0.32; fig. S4b). This is consistent with previous work showing the target is selected immediately after stimulus onset[22].

The cue decoding analysis replicates this result. In both monkeys, a decoder trained to infer the cue using only correct trials fails to generalize to swap trials (Monkey E: − 0.38 to − 0.20 difference in decoding performance, swap - correct, Monkey W: − 0.35 to − 0.23, fig. 3f). Further, the cue is encoded equally well during the first and second delays on correct trials (Monkey E: − 0.03 to 0.05 higher cue decoding performance on correct trials in delay 1 than 2, Monkey W: − 0.06 to 0.02; fig. 3g, left); the decrease in cue decoding only emerges on behavioral swap trials (Monkey E: 0.17 to 0.35 higher cue decoding performance on swap trials in delay 1 than 2, Monkey W: 0.13 to 0.24; fig. 3g, right). Overall, in the prospective task, we find evidence for what we term cue selection errors – where the cue is remembered correctly over the first delay period, but is corrupted when it is retrieved from working memory and used to interpret the incoming color information.

## 2 Discussion

Our results provide insight into the neural mechanisms underlying ‘swap’ errors. In summary, we found the neural correlates of swap errors emerged well before the animal’s decision. In both monkeys, we find strong evidence for errors that occur during selection from working memory. In one monkey, we found results that were consistent with both misbound color and mis-interpreted cue representations, indicating that misbinding errors and cue-related confusion may still explain some behavioral swap errors in some individuals. Furthermore, neither monkey showed evidence that swap errors could be explained by failing to encode or by forgetting one of the two stimuli after they were presented (see *Stimulus forgetting* in *Supplement* and fig. S3). Together, this work provides insight into the sources of errors in working memory. Previous work has posited that swap errors emerge due to the accumulation of errors during storage in working memory[27, 28] (though this effect is only pronounced for large set sizes). In contrast, we found evidence that even when storage in working memory is reliable, manipulating the contents of working memory can corrupt memories and lead to behavioral errors. Thus, our work provides fertile ground for a new theory of how information is manipulated in working memory, with implications for behaviors well beyond the tasks studied here.

## Acknowledgments

We are grateful to Stefano Fusi, Kirsten C.S. Adam, and Krithika Mohan for comments on earlier versions of this manuscript. We are also grateful to Allison Ong and Ciela Sophia Chavez-Gilbride for administrative support. This work was supported by the following grants: Simons Foundation 542983SPI (WJJ), Neuronex NSF 1707398 (MA and WJJ), Gatsby Charitable Foundation GAT3708 (MA and WJJ),the Swartz Foundation (MA and WJJ), NIMH R01MH115042 (TJB), and an NDSEG Fellowship (MP). We acknowledge computing resources from Columbia University’s Shared Research Computing Facility project, which is supported by NIH Research Facility Improvement Grant 1G20RR030893-01, and associated funds from the New York State Empire State Development, Division of Science Technology and Innovation (NYSTAR) Contract C090171, both awarded April 15, 2010.

## Author contributions

MP and TJB conceived of the experiments. MP performed the experiments. TJB supervised the experiments. MA and WJJ developed the statistical models and analytical approach. MA and WJJ performed the data analysis. MA and WJJ made the figures. MA, MP, TJB, and WJJ wrote and edited the paper.

## M1 Methods

### M1 Subjects

The experiments were performed in two adult Rhesus macaques (Macaca mulatta; 8-9 years old). Monkey E and Monkey W weighed 12.1 and 8.9 kg, respectively. All experimental procedures were approved by the Princeton University Institutional Animal Care and Use Committee and were in accordance with the policies and procedures of the National Institutes of Health.

### M2 Behavioral tasks

The behavioral task was implemented in Psychtoolbox and MATLAB (Mathworks) and it was displayed on a Dell U2413 LCD monitor, which the monkeys viewed at a distance of 58 cm. During the task, the animals were required to remember the two square stimuli presented at 45° clockwise (upper) and anti-clockwise (lower) from the horizontal meridian and with an eccentricity of 5 DVA. The colors were sampled from 64 uniformly spaced points on a photometrically isoluminant circle in CIELAB color space, centered at (*L* = 60, *a* = 6, *b* = 14) with a radius of 57 units. The two colors for each trial were sampled independently.

During an experimental session, the animal performed two different tasks in a blocked fashion. The first task is referred to as the retrospective task (retro). This tasks begins with a fixation period (500 ms) before two colored squares appeared on the screen for 500 ms. Then, after a delay period (500 ms or 1000 ms), a cue was presented in the fixation window (and subtending 2°) for 300 ms, which indicated to the animal which of the two colors should be reported. In the experiments, two distinct sets of cues were used: (1) lines oriented 45° clockwise or anti-clockwise from the horizontal meridian indicating the lower or upper position, respectively; and (2) a triangle or a circle, indicating the lower or upper position respectively. Next, after a second delay period (500 ms to 700 ms), a 2° wide color wheel centered on the fixation point appeared and the animal signalled their chosen color with a saccade to the corresponding location on the color wheel. The color wheel was randomly oriented on each trial to prevent motor planning or encoding of a spatial memory prior to the presentation of the wheel. The animals were rewarded according to the distance of their response from the target color (maximum of twelve drops, precise reward determined by a von Mises distribution centered at 0° error with a standard deviation of 22°). They were not rewarded if their response was greater than 40° or 60° from the target, for Monkey E and W respectively.

The second task is referred to as the prospective task (pro). In this task, after a fixation period (500 ms), the cue was shown (300 ms), followed by a delay period (200 ms to 600 ms). Then, the two colored squares were shown (500 ms) and after a second delay period (Monkey E: 1000 ms to 2000 ms; Monkey W: 1300 ms to 2000 ms), the animal made their response through a saccade onto the color wheel. The details of the color wheel and reward scheme were the same as for the retrospective task.

The two tasks were performed in randomly interleaved blocks. There were three blocks: prospective trials with the first cue set, retrospective trials with the first cue set, and retrospective trials with the second cue set. Block transitions occurred after 60 correct trials on the prospective task or 30 correct trials on either of the retrospective blocks (thus balancing the number of prospective and retrospective trials).

Throughout the whole experimental session, the eye position of the animal was monitored at 1 kHz through video-based eye tracker (SR Research). The monkeys had to maintain their fixation within 2° of a central fixation cross until the response period after the second delay. The trial was aborted and the monkey received a short time out if they moved their eye out of this window.

In addition to the two main experimental conditions that we analyzed here, the monkeys also both performed a version of the task where only a single color was shown. We do not show any primary analyses of these data, though we use it in the fitting of our neural mixture model (see *The neural mixture model* in *Methods*).

Monkey E completed 11,131 trials over 13 sessions and Monkey W completed 9,865 trials over 10 sessions.

### M3 Electrophysiological recordings

To immobilize the head, monkeys were implanted with a titanium headpost. Both monkeys were also implanted with two titanium recording chambers, which were positioned using 3D models of the skull to allow for recordings from LPFC, FEF, partial cortex, and V4.

Neural activity was recorded via up to 80 simultaneously inserted epoxy-coated tungsten electrodes (FHC). The electrodes were allowed to settle for 2-3 hours before recordings. Broadband (30 kHz) activity was recorded from each electrode (Blackrock Microsystems).

### M4 The behavioral model

To classify responses as correct, a swap, or a guess, we modeled the responses on each trial as a mixture of three different possible response types: a correct response, swap response, and guess response. In particular, we fit the model:

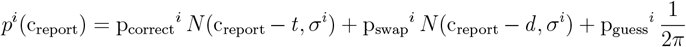

where 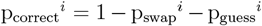 and *N* (*μ, σ*) is the probability distribution function of a normal distribution with mean *μ* and standard deviation *σ*. The index *i* related to experimental sessions. We fit the model in a hierarchical manner within each monkey, where the probability parameters were modeled as arising from a distribution across distinct sessions. In particular, we reformulated the problem using the transformation:

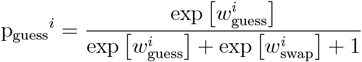

and similarly for 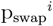 while 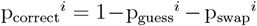. Then, we modeled 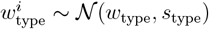 for guesses and swaps. This hierarchical approach provides the benefit of partial pooling, where the different error rates on each individual session are not assumed to be exactly the same nor are they assumed to be completely independent[29]. In total, our behavioral model had 3 parameters for each session (p_swap_, p_guess_, *σ*) and 4 parameters at the level of each monkey (*w*_swap_, *w*_guess_, *s*_swap_, *s*_guess_. The monkey-level mean parameters were given normal priors with mean 0 and standard deviation 1, while the standard deviation parameters were all given half-normal priors with mean 1 standard deviation 3. We obtained qualitatively similar results when fitting the models at the per session level with a uniform Dirichlet prior for the probability parameters.

We also implemented an inverse model from which we could sample from a posterior predictive distribution of response errors. In particular,

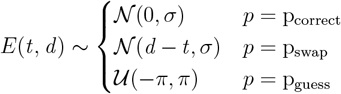

where all values are treated as periodic between −*π* and *π*.

Finally, for each trial, we also computed the posterior probability that the response arose from each of the three categories that we considered, according to Bayes rule:

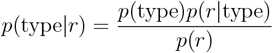

#### M4.1 Fitting procedure

The models were fit using Hamiltonian Monte Carlo (HMC)[30] with No-U-Turn Sampling (NUTS)[31] implemented in the probabilistic programming language Stan[32] which we interacted with via the pystan[33] package. For each model, we ran 4 chains and for 500 warmup iterations and 500 samples. We verified that the model had converged via the 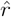 scores for each parameter, which quantifies how well the different chains have converged to the same distribution. In particular, 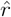 is 1 if the chains have converged to the same distribution and *>* 1 otherwise. Each parameter from the fit models reported here had 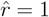. We also evaluated other metrics of fit, which yielded no warnings.

### M5 Hypothesized error types

Across our two primary epochs of interest, we attempt to explain swap errors (that is, the behavioral swap response) in four different ways.

#### M5.1 Misbinding errors

In this case, either when the stimuli are initially encoded or while they are being stored in working memory over the first delay period, the color and location associations are misbound to each other – such that a green color in the upper location and a blue color in the lower location are represented as if they were a blue color in the upper location and green color in the lower location. We incorporated this possibility into our mixture model as described below (*The mixture model* in *Methods*). This form of error has been theorized for decades, both in psychology[14–16] and in neuroscience[17, 18, 34].

#### M5.2 Cue interpretation errors

In this case, the meaning of the cue is misinterpreted on certain trials (e.g., a cue indicating the animal should report the color at the lower position is interpreted to mean that the animal should report the color at the upper position), which leads to a behavioral swap response. We hypothesize that this difference would manifest in the neural representation of the cue. In particular, we expect that the neural representation would reflect the animal’s belief about what the cue was – and on swap trials, this would look like a representation of the incorrect cue.

#### M5.3 Selection errors

In both tasks, the animal first remembers one piece of information, then has to combine that piece of information with a second piece of information. In particular, in the retrospective task, the animal first sees two colors and must remember both the colors and their positions. Then, the animal is shown a cue. To solve the task, the animal has to use that cue to select one of the two remembered colors (based on its position) for eventual report. For selection errors, we hypothesize that it is this selection of information that was previously (and successfully) stored in working memory that gives rise to swap errors, by leading to an error in encoding when it is selected.

#### M5.4 Forgetting

One potential explanation for swap errors is that, just after the stimuli have been presented, the animal simply forgets one of the two stimuli. Then, when presented with the response wheel, the animal simply reports the color that they remember even though they know that it may not be correct. We had difficulty incorporating this forgetting possibility into our mixture model, so we instead worked to evaluate the number of stimuli encoded during the second delay period (either 1 or 2), using a subset of trials on which a single stimulus was presented. Here, we show that swap errors are not associated with representations that look like those on trials where a single stimulus had been presented (for more detail see *Stimulus forgetting* in *Supplement*), which is inconsistent with the educated guessing explanation for swap errors proposed in [20].

### M6 Electrophysiological data preprocessing and exclusion

We include both single and multi-units in our analyses. We exclude units that have zero firing rate for a contiguous 50 trial block in the session from analysis. After this exclusion, we bin the firing rates into 500 ms bins (except in the case of the first delay period on prospective trials, since the delay period is only 200 ms, we use only 200 ms bins) and then z-score each neuron individually before performing principal components analysis and retaining 95 % of the variance. In the cross-validated analyses both of these transformations are fit to only training-set trials, then applied to the test-set trials.

### M7 The neural mixture model

To model the neural data, we also take a mixture modeling approach. The model can be understood as having two parts. The first part models the responses to the two presented colors and (in delay 2) to the cue. The second part of the model, models to dependence of the responses on the eventual behavioral response – i.e., a correct response, swap error, or guess. We will describe these two components in sequence.

#### M7.1 Additional preprocessing

For this model, we add two additional preprocessing steps, which do not qualitatively effect our conclusions but which do improve the convergence times of our models. Even after our exclusion of neurons with consistently zero firing rates for 50 trials in a row during the session, many neurons in the population have zero firing rate for some contiguous portion of the session. To ameliorate this, we impute the firing rate of those neurons for their stretch of contiguously zero firing based on the 5 nearest neighbor trials across the population (KNNImputer in sklearn[35]).

We also z-score the firing of each unit after the PCA transform, since we use a uniform prior variance for the firing of each unit in the statistical model.

#### M7.2 The color and cue representations

We model the color representations using two sets of periodic splines. In particular, for a particular color *c* ∈ [0, 2*π*], we map it into a spline representation with the function **f** (·), which moves from a single number to a *K*-dimensional representation where *K* is the number of “knots” in the spline representation. We also control the smoothness of the splines with an order parameter. We performed all of our analyses with *K* = 4, 5, 6 and order = 1, 2. The results did not qualitatively change for any of these choices. We report results from models with splines chosen to have *K* = 5 and order = 1.

Once we have the spline representation of both the upper and lower (target and distractor) colors in delay 1 (delay 2), then we model the neural responses *r* as a linear model of both colors:

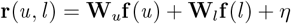

where **W**_*u*_ and **W**_*l*_ are fit parameters, *u* and *l* are the upper and lower colors (for delay 1), and *η* ∼ 𝒩 (0, *σ*). Thus, the parameters fit for this part of the model are **W**_*u*_ and **W**_*l*_, which are each *N* × *K* matrices and *σ*, which is a *N* × 1 vector (we do not fit a full covariance matrix), where *N* is the number of neurons (or the number of dimensions after PCA) and *K* is again the number of knots in the spline basis.

To model the cue, we simply add a binary variable *c* which indicates whether the upper color (*c* = 1) or lower color (*c* = 0) are cued for report:

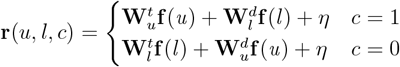

Now there are four matrix parameters, for the upper and lower color when they are target or distractor. For example, 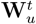 is the representation of the upper color when it is the target, while ^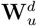^ is its representation when it is the distractor. This allows for a flexible arrangement of the color represenations.

#### M7.3 The mixture model

We include a mixture component with the model described above, where for a given trial during delay 1, we fit,

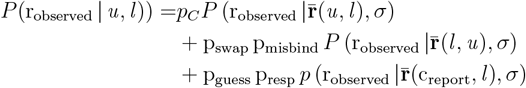

where *p*_*C*_ = p_correct_ + p_swap_(1−p_misbind_)+p_guess_(1−p_resp_) is the accumulated probability of the correct representation, p_correct_, p_swap_, p_guess_ are taken from the behavioral model and are not fit, p_misbind_ and p_resp_ are free scalar parameters that represent the likelihood that a given response looks like either a misbinding error (*u* and *l* are swapped) or a particular version of a guess, in which the representation of the eventually reported color looks more like what the animal eventually reports (c_report_) than the actual color (as in this example, *u*). This part of the full model adds two parameters to the total set of parameters fit for the neural mixture model. Importantly, all of the parameters are fit simultaneously, rather than in two stages.

The model for delay 2 in the retrospective task is similar, but has another possibility for swap errors that relates to the encoding of the cue. Here, we take an example trial where the cue indicates that the upper stimulus is the target,

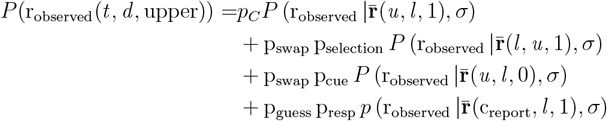

where *p*_*C*_ = p_correct_ + p_swap_(1 − p_selection_ p_cue_) + p_guess_(1 − p_resp_) is the accumulated probability of the correct representation.

The model for delay 2 in the prospective task is identical, but the interpretation of the mixture parameters is different. In particular, p_selection_ from the retrospective task is analogous to p_misbind_ and p_cue_ is analogous to p_selection_, as discussed in the main text.

#### M7.4 Hierarchical structure for the delay 2 models

To improve the quality of our parameter estimates, we adopt a hierarchical stucture for our delay 2 model, since the set of predictions that we test are the same across both the retrospective and prospective tasks. In this structure, we fit

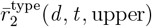

for each type ∈{retro, pro, single} and where each of the type-specific *W* and *z* are assumed to be drawn from a normal distribution with a mean and standard deviation that is fit across all the types. These combined mean and standard deviation parameters are given normal and half-normal priors, respectively – both with standard deviation 10. As with the behavioral model, this allows us to partially pool information across the different task conditions, where the representation of color is not assumed to be exactly the same across tasks – but nor is it assumed to be completely independent.

#### M7.5 Fitting procedure

As for the behavioral model, we fit this model using HMC in Stan. For each model, we ran 4 chains and for 500 warmup iterations and 500 samples. We verified that the model had converged via the 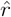 scores for each parameter, which quantifies how well the different chains have converged to the same distribution. In particular, 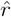 is 1 if the chains have converged to the same distribution and *>* 1 otherwise. Each parameter from the fit models reported here had 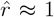. We also evaluated other metrics of fit, which yielded no warnings.

### M8 Cue decoding analysis

For the cue decoding analysis, we began by splitting the trials into likely correct trials (p_correct_ *>* .6) and likely swap trials (p_swap_ *>* .4). Then, we train a linear classifier to decode the value of the cue on the likely correct trials. The reported decoding performance is the cross-validated performance of the decoder on those trials. Next, we evaluate the performance of the same decoder to all of the likely swap trials.

Thus, if likely correct and likely swap trials have similar representations of the cue, we expect the decoder to generalize (as for the first delay period of the prospective task). However, if the representations differ, then the decoder will not generalize (as for the second delay period of the prospective task).

### M9 Cross-validated neural model

To evaluate how the observed effects change over time as well as to replicate our findings from the Bayesian neural mixture model with more familiar techniques, we develop a fully cross-validated version of the mixture model analysis. Here, we again divide the trials into likely correct (p_correct_ *>* .3) and likely swap (p_swap_ *>* .3) trials. We also exclude any trials where the target and distractor colors are closer than 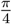 radians apart. Then, we fit the model described in section M7.2 as a standard Ridge regression using only likely correct trials in a leave-one-out scheme. Thus, for that particular fit, we have one likely correct trial that has not been used to fit the model as well as several likely swap trials. For each of those trials, we use the model to construct our usual hypothesis space (e.g., the prototypical correct and spatially misbound representations from the first delay period of the retrospective task) and we evaluate the projection of the held out trials on the dimension connecting these different prototypes. In particular, for each each held out trial *r* and two prototypes 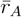 and 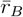, we compute,

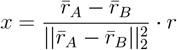

and *x* will be close to zero if *r* is close to *r*_*B*_ while it will be close to one if *r* is close to *r*_*A*_. Thus, to evaluate the evidence for swap errors, we compare the average value of *x* taken across likely correct with the average value taken across likely swap trials.

In this analysis, we consider all of the same prototypes that we consider in the mixture model.

## S1 Supplement

### S1 Region-dropping analysis

To understand whether the different regions recorded here contribute differently to the neural correlates of swap errors, we have performed a region dropping analysis using the cross-validated timecourse model described in the main text. To evaluate the contribution of a particular region, we fit the model as described in the main text and calculate the difference between correct and swap traces. Then, we evaluate how this difference changes, when we exclude either all of the neurons from a single region or an equivalently sized set of random neurons. We choose 100 random, equally sized subsets to remove. If a particular region is more important for the neural correlates of swap errors than average, then this difference will be positive (in the figure, we refer to this as the relative importance). If a region is less important than average, then this difference in effects will be negative.

Overall, this analysis finds only inconsistent regional effects (fig. S1). There is some evidence in Monkey W and specifically in the prospective task (fig. S1, right) that the neural correlates of swap errors emerge first in V4/PIT (i.e., evidence for swap errors during delay 1) and then transition into PFC (i.e., evidence for swap errors during delay 2) – but the second monkey does not show a reliable effect in the same direction. This analysis supports the idea that working memory representations are broadly distributed across different brain regions.

### S2 Analysis of guess responses

The neural mixture model finds high probability that guess trials resemble representations of the eventually reported stimulus in both the first (fig. S2b) and second (fig. S2c delay periods of the retrospective task. However, the cue identity appears to be reliably encoded in the second delay period, indicating that the animal does not forget all experimental variables on guess trials (fig. S2d). The same pattern of results holds in the prospective task (fig. S2e, f, g), though there is a less reliable representation on the cue on likely guess trials in one of the two monkeys.

### S3 Stimulus forgetting

We investigate whether the the neural activity on swap trials is consistent with a representation of just one rather than two total stimuli. This addresses the explanation for swap errors proposed in [1].

To do this, we train a linear decoder to discriminate between trials from the retrospective (fig. S3a) or prospective (fig. S3b) tasks (both in the second delay period) and trials on which only a single color was shown. Then, we test evaluate the generalization performance of that decoder on likely swap trials. If behavioral swap errors result from only a single stimulus being remembered and the animal simply reporting that remembered stimulus, then we expect that the decoder will fail to generalize. However, we find that the decoder performs just as well on swap trials as on correct trials

**Figure S1:**
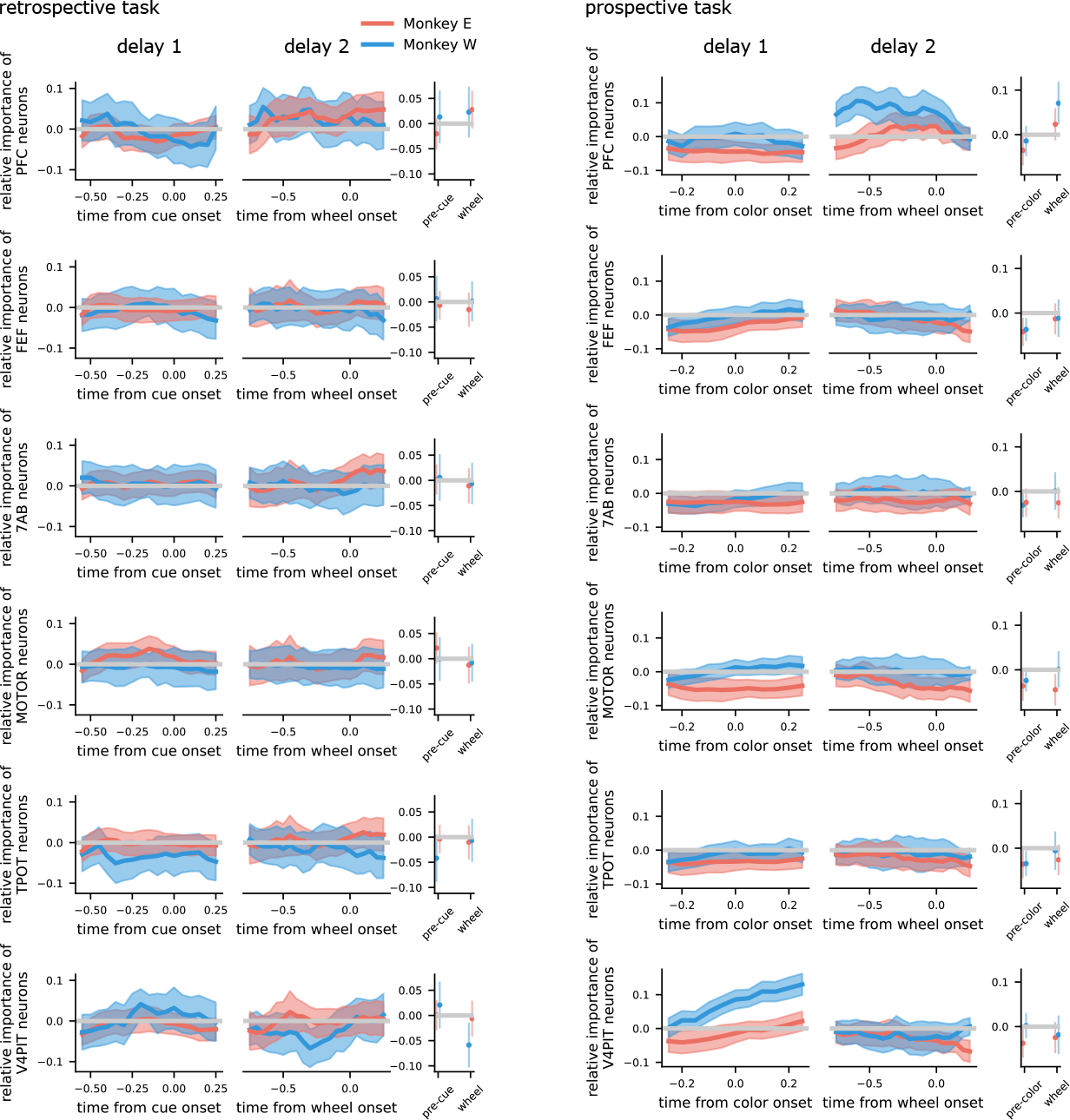
The unique contribution of different brain regions to the neural correlates of swap errors. (left) The region dropping analysis applied to delay 1 and delay 2 of the retrospective task. The right column shows the relative important at the specific times reported in the main text. There are not consistent effects across monkeys. (right) The region dropping analysis applied to delay 1 and delay 2 of the prospective task. The layout is the same as for the retrospective task and there are no consistent effects across monkeys.

---

**Figure S2:**
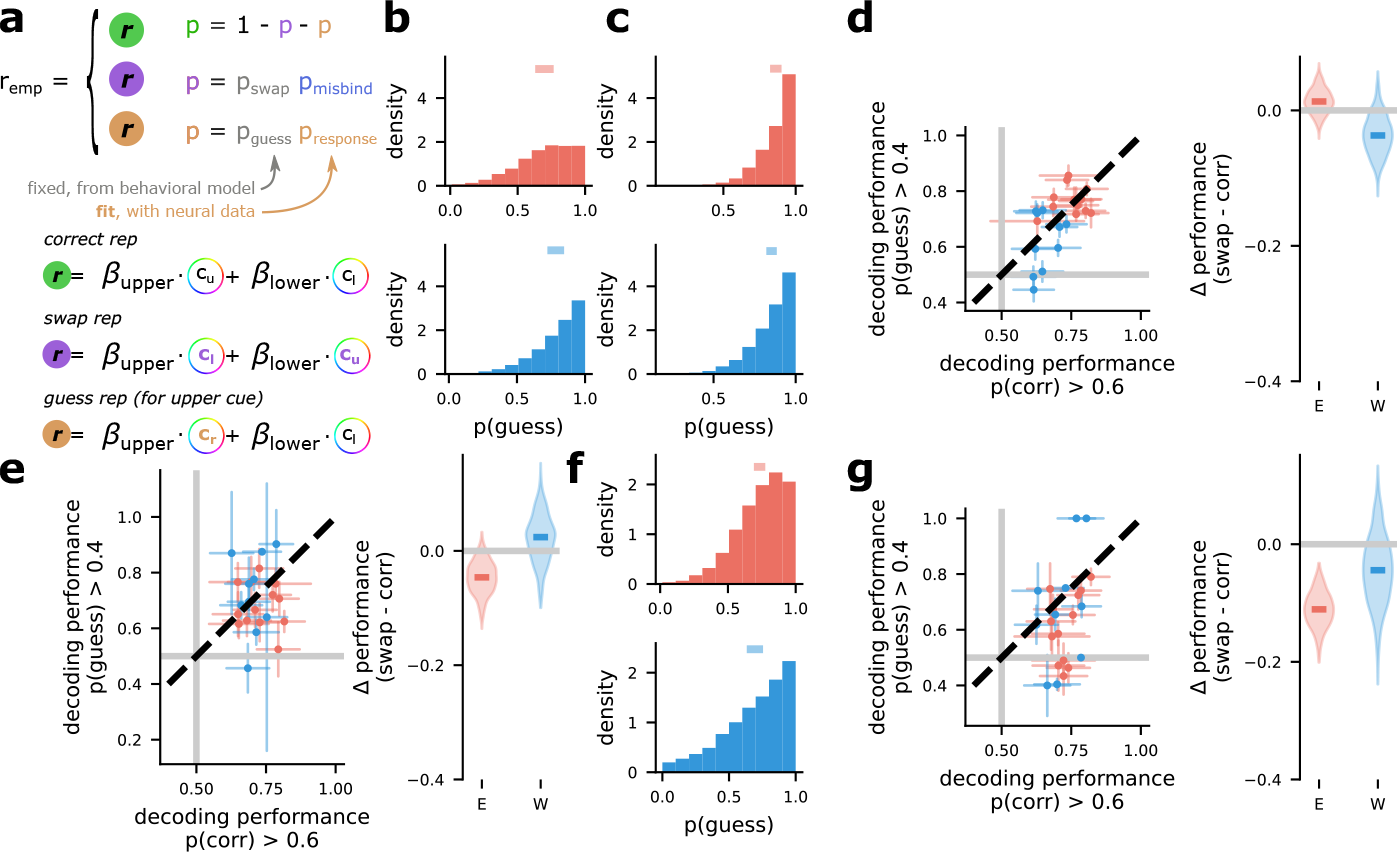
The neural correlates of guess responses in retrospective and prospective working memory tasks. **a** Schematic of the mixture modeling approach used to express guess responses. Here, we assume that the neural activity on guess trials will represent the eventually reported color (*c*_*r*_), even when that color is neither the target nor the distractor. **b** Average (line) and aggregate posterior (distribution) across sessions for the *p*_response_ parameter shown in **a** during delay 1 of the retrospective task. **c** The same as **b** but for delay 2 of the retrospective task. **d** The decoder generalization analysis applied to delay 2 of the retrospective task, but instead of contrasting likely correct and likely swap trials the analysis contrasts likely correct and likely guess trials. **e** The same as **d** but applied to delay 1 of the prospective task. **f** The same as **c** but applied to delay 2 of the prospective task. **g** The same as **d** but applied to delay 2 of the prospective task.

**Figure S3:**
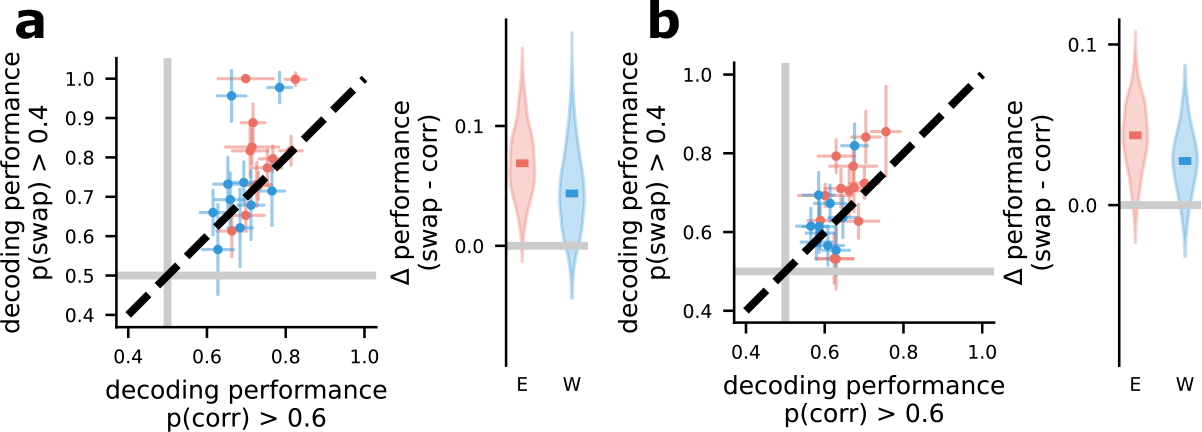
Swap errors are not associated with the encoding of a different number of stimuli in working memory. **a** The performance of a decoder trained to decode the number of stimuli shown during the pre-response wheel period (delay 2) on the retrospective task. (left) The x-axis shows decoding performance on the likely correct trials, while the y-axis shows the same decoder generalized to likely swap trials. (right) Average difference for both monkeys. **b** The same analysis as **a** but for the prospective task.

**Figure S4:**
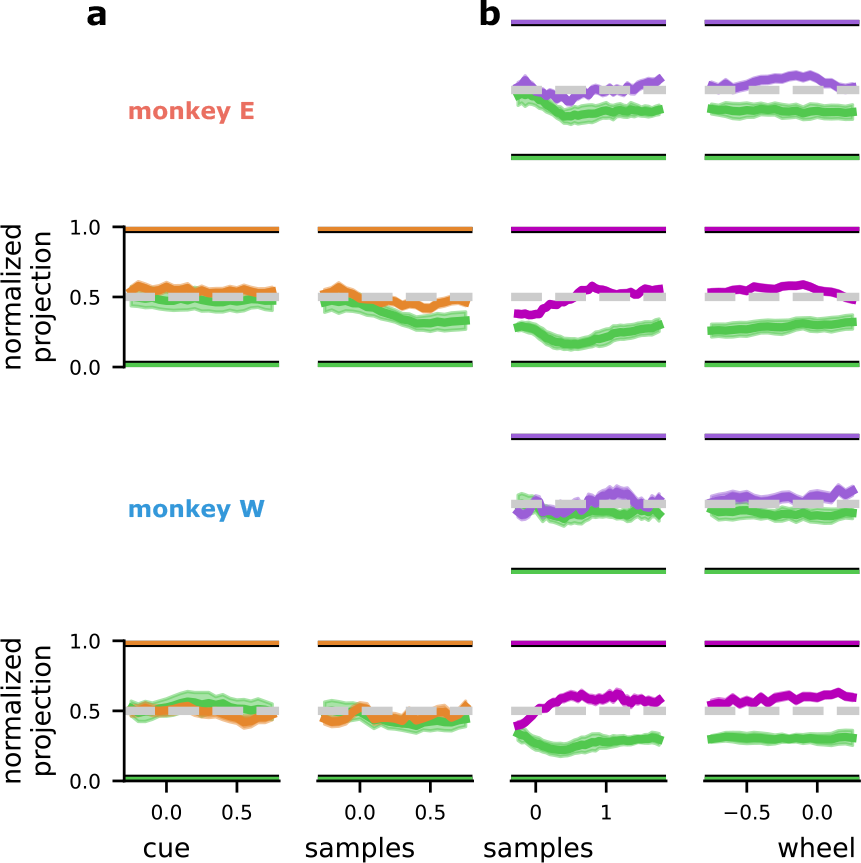
The timecourse of evidence for neural correlates of swap errors in the prospective task. **a** Evidence for cue interpretation errors in monkey E (top) and monkey W (top). **b** Evidence for misbinding (first and third rows) and cue selection (second and fourth rows) in both monkeys (top and bottom blocks).

